# Bacterial Engineered Living Materials modulate Mechanosignaling in Mammalian Cells

**DOI:** 10.1101/2024.10.29.620857

**Authors:** Katharina Ostmann, Geisler Muñoz-Guamuro, Jan Becker, Miguel Baños, Sairam Saikumar, Roland Bennewitz, Cao Nguyen Duong, Shrikrishnan Sankaran, Wilfried Weber

**Affiliations:** INM – Leibniz Institute for New Materials, Campus D2 2, 66123 Saarbrücken, Germany; Signalling Research Centres BIOSS and CIBSS, University of Freiburg, Schänzlestraße 18, 79104 Freiburg, Germany; Spemann Graduate School of Biology and Medicine (SGBM), University of Freiburg, Albertstraße 21a, 79104 Freiburg, Germany; Faculty of Biology, University of Freiburg, Schänzlestraße 1, 79104 Freiburg, Germany; Saarland University, Department of Materials Science and Engineering, Campus D2 2, 66123 Saarbrücken, Germany; Signalling Research Centres BIOSS and CIBSS, Faculty of Biology, and Spemann Graduate School of Biology and Medicine (SGBM), University of Freiburg, 79104 Freiburg, Germany

**Keywords:** Biofilm, Curli, extracellular matrix, living therapeutic material, wound dressing, YAP/TAZ

## Abstract

Engineered living materials (ELMs) are gaining momentum for biomedical applications as self-replenishing drug depots, smart wound dressings, or as wearable sensors. Current studies on ELM-host interaction are mainly limited to the exchange of biochemical cues between ELMs and surrounding cells and tissues. Here we show that the genetically programmed mechanical properties of ELMs modulate mechanosignaling pathways in mammalian cells cultivated onto the living materials. To this aim, we genetically modulated curli fiber production in *E. coli* and analyzed the impact on the mechanical properties of the resulting ELMs. The living materials were used as matrix for the cultivation of mammalian cells engineered with a fluorescent reporter to indicate the activation of the mechano-responsive Hippo signaling pathway. We demonstrate that different genetically programmed ELM compositions translated into differential regulation of mechanosignaling in mammalian cells. These findings provide the perspective of using ELMs as extracellular matrix with genetically programmable mechanics for mammalian cells while also highlighting the need to consider the mechanical properties of therapeutic ELMs when assessing interaction with surrounding tissues.

ToC figure

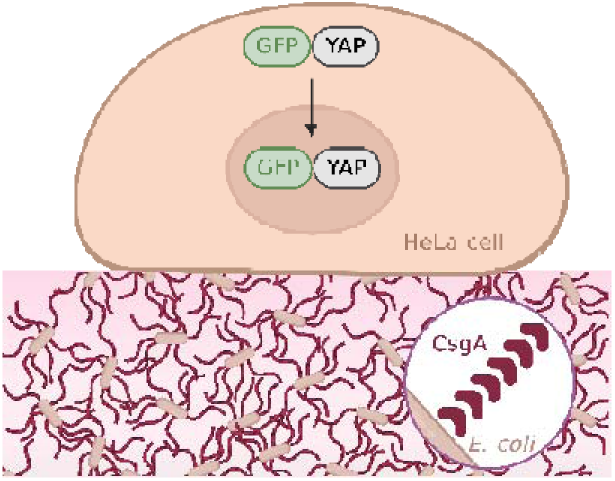

## Introduction

Engineered living materials (ELMs) are an emerging class of materials that contain living cells, which either form their own matrix around themselves or are incorporated within non-living materials.[1] Those living cells can be programmed to react to environmental stimuli, self-regenerate and self-assemble. Based on the functionality of the living cells, ELMs are being developed for a variety of applications such as sensors, textiles, structural materials, or living electronics .[2–5] Recently, there has been a rise in the development and optimization of ELMs for biomedical applications. These include, for example, the development of wearable biosensors, skin patches for wound healing, smart wound dressings, drug delivery systems as well as stem cell and tissue engineering.[6,7,16,8–15] A large variety of organisms have been utilized for such applications including bacteria, fungi, or algae.[17–20] To shield the organisms from the host, they are often encapsulated in polymeric matrices composed of agarose, Pluronic F-127 or alginate.[16,21,22] For example, Lufton et al. developed antifungal skin patches incorporating *Bacillus subtilis* in a Pluronic F-127 gel.[7] By varying the composition of this gel, mechanical properties can be adjusted which is important for the integration of the material with host tissues.

In contrast to such hybrid ELMs, native- or bio-ELMs consist of cells that make their own matrices. This category of ELMs allows genetic encoding of both cellular functions and material properties. A prominent approach to make such ELMs involves *Escherichia coli* producing curli fiber-based biofilms. These are protein fibers that have been modified to create self-produced matrices with novel (therapeutic) properties. The fibers consist primarily of the protein CsgA that is secreted by the bacteria and nucleates at the bacterial surface supported by the protein CsgB, which leads to the formation of extracellular fibers.[23] The fibers possess the advantage of being easily produced, genetically modifiable and biodegradable.[24] For example, towards the development of therapeutic ELM, probiotic *E. coli* have been engineered to produce a modified variant of CsgA fused to a human trefoil factor (TFF) peptide that promotes epithelial barrier restoration in the gut for treating inflamed bowel tissues.[19]

Notably, the introduction of foreign materials into the body can cause a variety of host reactions. Thus, different strategies to ensure biocompatibility of ELMs have been implemented, for example, by using organisms with Generally Recognized as Safe (GRAS) status, such as *Lactococcus lactis* or *Bacillus subtilis* or by encapsulating the bacteria in biocompatible materials.[25–27] These strategies to enhance biocompatibility of ELMs focus mainly on mitigating biochemical effects such as the production of reactive molecules. One factor which has largely been neglected in such analysis is the direct interaction of ELMs with surrounding cells in the context of mechanosignaling. Mechanosignaling is an important process in shaping mammalian cell fate and function[28,29] by controlling important processes such as cell development, differentiation, adhesion, migration, proliferation, and ECM generation.[30] A dysregulation in the mechanical properties of the cellular environment is associated with pathological processes including fibrosis, tumorigenesis, and cancer immunotherapy resistance.[30] Different mechanisms are responsible for the transduction of mechanical cues into a cellular response. This includes cytoskeleton remodeling and activation or deactivation of biochemical pathways. Two key protein effectors involved in mechanotransduction are the transcription factors YAP (Yes-associated protein) and TAZ (Transcriptional coactivator with PDZ-binding motif).[31] They are regulated by a variety of signaling pathways, such as Hippo or Wnt, which are crucial in regulating tissue homeostasis and organ size.[32] It has been shown that changes in the mechanical properties of the cellular environment, for instance increased stiffness, can activate YAP/TAZ. This activation leads to their translocation to the nucleus where they regulate the expression of various genes, with functions including the stimulation of cell proliferation by reprogramming cell metabolism via transcriptional regulation of involved enzymes and biosynthesis machineries.[33]

In this study, we develop a method for modulating the mechanical properties of curli-based ELMs by engineering the CsgA content. We demonstrate that the engineered ELMs triggered differential YAP localization upon co-culture with HeLa cells. These studies suggest that considering (genetically encoded) stiffness of ELMs may be important in avoiding interference with cellular signaling cascades. It further opens the perspective of using ELMs as extracellular matrix for tissue engineering with genetically programmable mechanical properties for steering cell fate and function.

## Results and Discussion

In order to study the mechanical influence of curli fiber-based ELMs on mammalian cells, we developed two curli-producing *E. coli* biofilms with different stiffnesses, and a mammalian reporter cell line based on hYAP1 (human Yes associated protein 1) (**Figure 1**).

**Figure 1.**
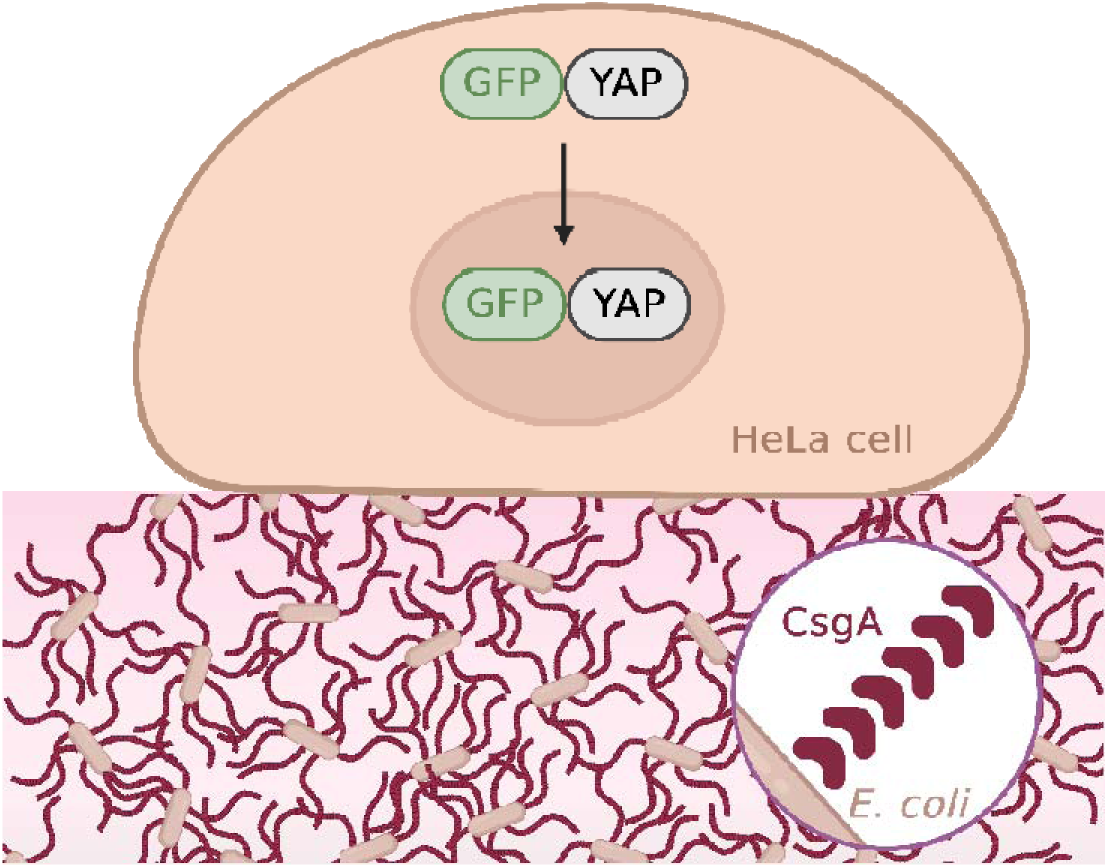
Schematic overview. *E. coli* biofilms, producing different levels of the extracellular protein CsgA exhibit different stiffness. We analyzed the impact of these ELMs on mechanosignaling by employing a HeLa reporter cell line based on an eGFP-hYAP1 fusion protein.

First, we developed and characterized the two biofilms with different storage moduli. We used the *E. coli* PHL628 strain, a K12 MG1655 derivative with an *ompR234* mutation, which leads to a higher expression of curli fibers (hereafter called CsgA_High_).[34,35] Hypothesizing that lower amounts of curli fibers lead to biofilms with a less dense fiber network and different mechanical properties, we engineered a CsgA knockout strain (PHL628 Δ*CsgA*) with a plasmid encoding CsgA and mCherry under the control of the Trc promoter leading to 4.8-fold lower levels of curli fibers (**Figure S1**).[35] mCherry is used for visual confirmation of transformation and for discrimination between the engineered biofilm (hereafter called CsgA_Low_) and the CsgA_High_ biofilm. Under natural light, CsgA_High_ appears as a yellowish biofilm and CsgA_Low_, due to mCherry expression, as a pink biofilm (**Figure 2A**). In Figure 2A the curli fibers were stained with Congo red.[36] Accordingly, the CsgA_High_ ELM shows an intense dark red color compared to the knock-out strain.

**Figure 2.**
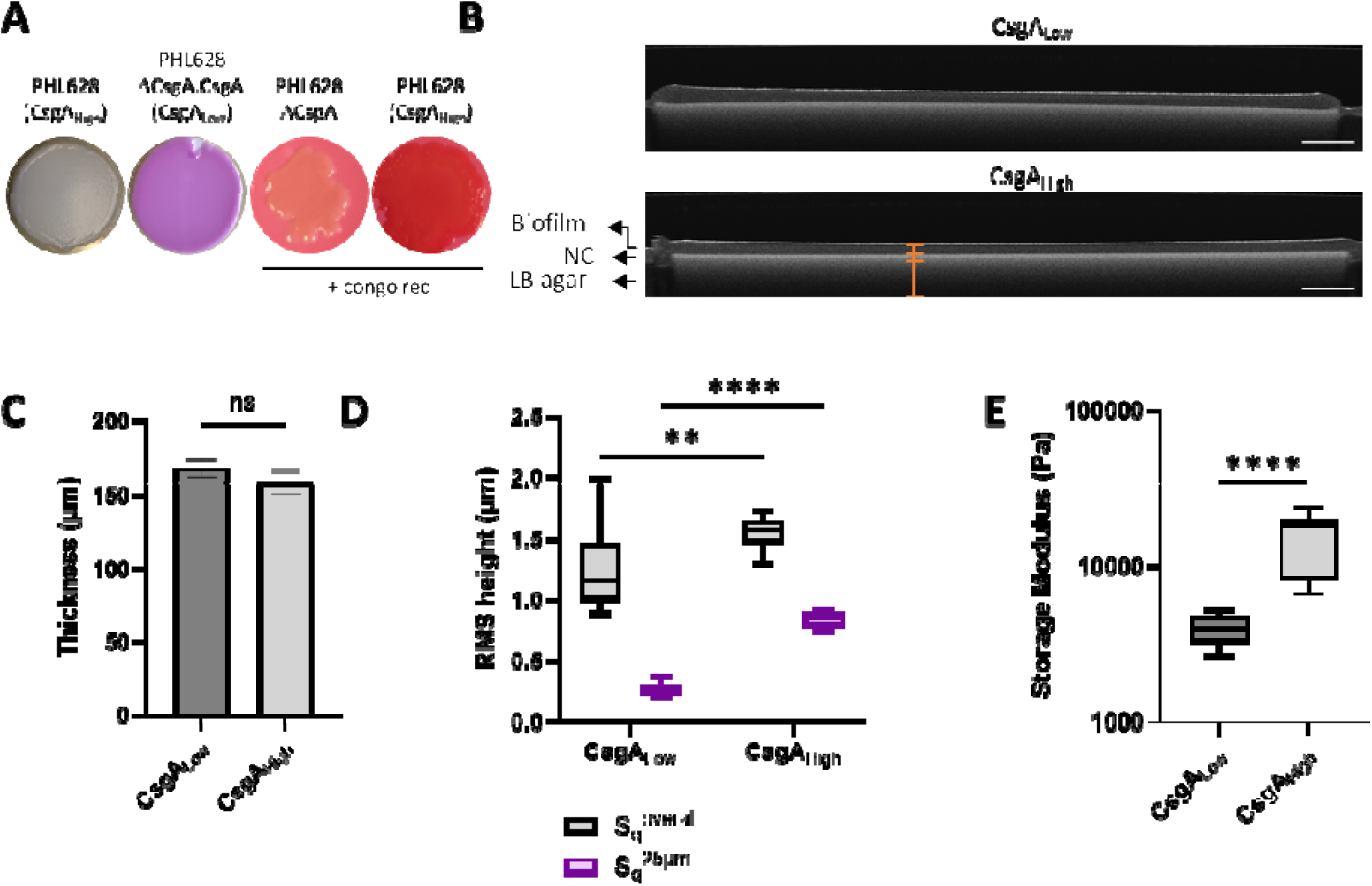
Characterization of biofilms. A Images of Congo red stained biofilms CsgA_High_ and CsgA_Low_. B Representative OCT cross-sectional image of biofilms. scale bar: 1000 µm. C Thickness of biofilms as determined from the OCT cross-sectional images. A refractive index of 1.37 was used in the calculations of the thickness for both biofilms (Figure S2). D Surface roughness of biofilms analyzed using 3D confocal microscopy. Two roughness parameters were calculated: The overall roughness S_q_^overall^ and the roughness S_q_^25µm^ at the scale of a single mammalian cell. E Rheology analyses of the CsgA_High_ and CsgA_Low_ biofilms. Data in C-E are representative from n = 3 replicates. Boxes represent median and upper and lower quartiles, whiskers represent min and max values. Mann-Whitney U test, \*\*\*\**P* ≤ 0.0001, \*\**P* ≤ 0.01, ns *P* > 0.05

For comprehensive characterization, we further determined the thickness of the biofilms from optical coherence tomography (OCT) cross-sectional images (**Figure 2B**). A refractive index of 1.37 was applied in the thickness calculations for both biofilms (**Figure S2**). The average thicknesses for CsgA_High_ (158.6 ± 8.0 µm) and CsgA_Low_ (167.8 ± 6.4) biofilms did not show a statistically significant difference (*P* > 0.05) (**Figure 2C**).

To the naked eye, the two ELMs appear to have different surfaces, with CsgA_High_ seeming to have a more structured surface than CsgA_Low_. To corroborate this observation, we further investigated the roughness of those two biofilms by using a 3D confocal microscope. To this end we determined two roughness parameters: the root mean square height (RMS height S_q_^overall^ at large scale and RMS height S_q_^25µm^ at the scale of a single mammalian cell. The S_q_ parameter is often used to evaluate the overall roughness of materials. It represents the RMS height deviation of each point of a surface profile measured from the mean plane. Our results show that the CsgA_High_ exhibits a significantly higher S_q_^overall^ than CsgA_Low_, namely 1.55 ± 0.13 and 1.25 ± 0.32 µm, respectively. While these values of S_q_^overall^ already provide an overall assessment of surface roughness, S_q_^25µm^ offers additional insight into the surface roughness at the length scale of a single mammalian cell. Our results reveal that CsgA_High_ has also a higher S_q_^25µm^ than CsgA_Low_, being 0.84 ± 0.06 µm and 0.27 ± 0.05 µm, respectively (**Figure 2D**). Consistently, topography heat maps obtained from the 3D confocal analysis show a higher number of peaks and grooves at this scale for CsgA_High_ than for CsgA_Low_ (**Figure S3**). In summary, the results reveal that CsgA_High_ has a significantly rougher surface than CsgA_Low_. This difference in roughness between the two biofilms may be attributed to the distinct composition of their extracellular matrices.

To analyze the stiffness of the biofilms, a rheometer was used to perform small oscillatory amplitude shear measurements and record relevant biofilm mechanical properties.

Rheological analysis showed a storage modulus G’ of 4.0 ± 0.8 kPa for CsgA_Low_ and of 15.7 ± 5.9 kPa for CsgA_High_ (**Figure 2E**, **Figure S4**).

In order to co-cultivate the mammalian reporter cells and the biofilms, the bacteria needed to be inhibited, because otherwise the bacteria would rapidly consume nutrients and acidify the medium by producing acetic acid. Such acidification can be observed by a change of color from red to yellow of the cell culture medium containing phenol red as pH indicator.[37,38] As a first try, we tested different bacteriostatic antibiotics, erythromycin, tetracycline, and sulfamethoxazole. To this end, CsgA_High_ biofilms were incubated in cell culture medium containing the pH-sensitive phenol red dye and different antibiotics in a range of concentrations (**Figure 3A**). It can be observed, that after 2 h of incubation, the medium turned yellow for all samples as quantified by measuring the absorbance at 560 nm (red state) and 430 nm (yellow state) (**Figure 3B, 3C, S5**). [39] These results align with previous research indicating that biofilms are resistant to a range of antibiotics.[40]

**Figure 3.**
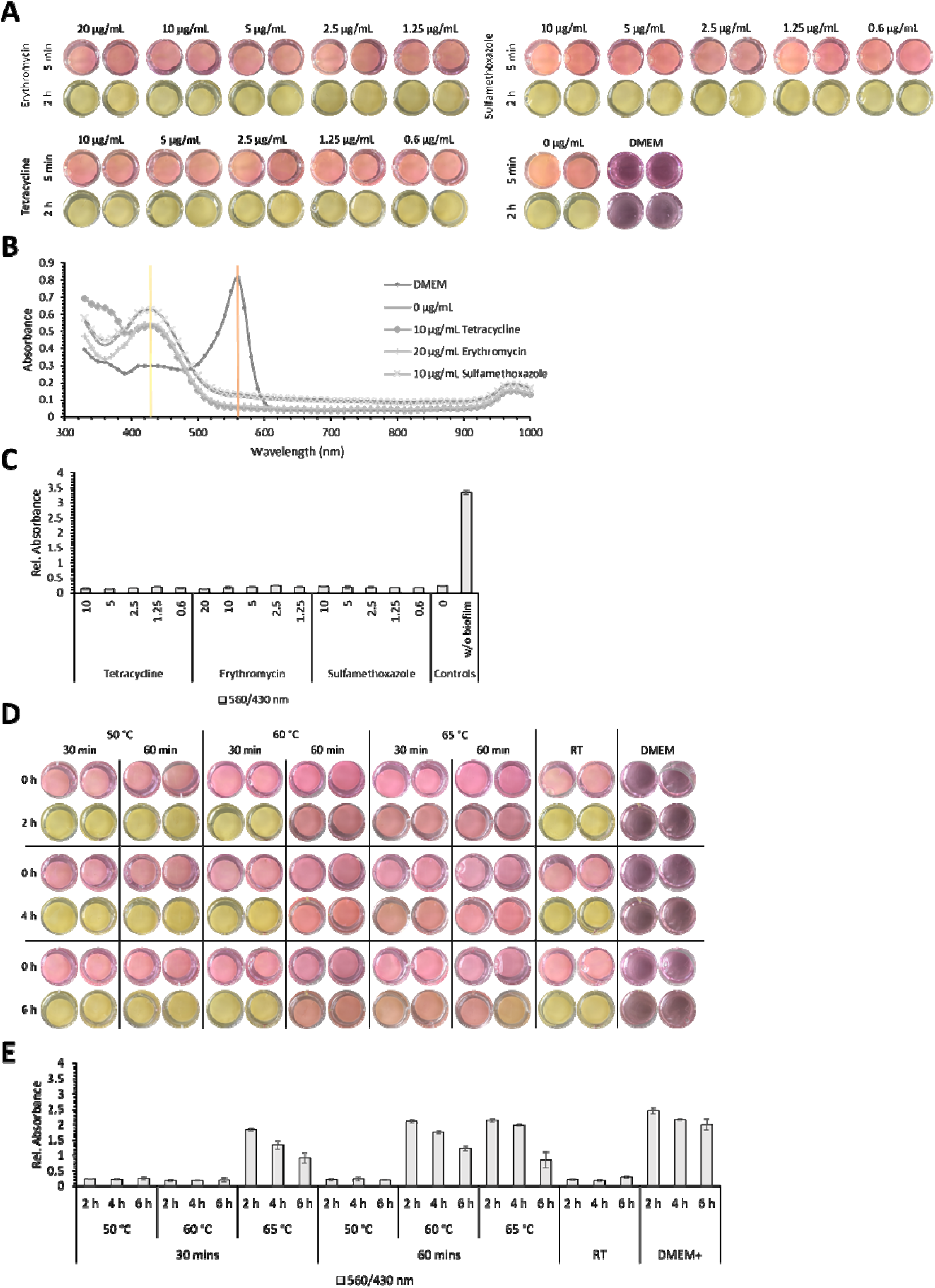
Co-cultivation of HeLa cells and biofilms. A CsgA_High_ (PHL628) biofilms were cultivated on nitrocellulose membranes placed on LB Agar plates for 4 days at room temperature. Afterwards, the biofilms were placed in 24-well plates and incubated in cell culture medium containing the indicated concentrations of antibiotics. After 2 h of incubation, images of the wells were taken and the supernatant was subsequently used to measure absorbance spectra. B Exemplary absorbance spectra of the medium after 2 h incubation.

As a next step, we evaluated heat treatment to inactivate the bacteria. For this, CsgA_High_ biofilms were first treated using different temperatures for different time durations (50-65 °C for 30-60 minutes). Following this heat treatment, medium color was assessed after 2 h, 4 h and 6 h of incubation in phenol red-containing cell culture medium (**Figure 3D, 3E, S6**).

Since the treatment at 65 °C for 60 min showed robust prevention of a change in color, this condition was used for the subsequent co-cultivation experiments.

Indicated as yellow and red lines are the absorbance peaks at 430 nm and 560 nm, respectively. C Representation of the relative absorbance at the peaks indicated before (560 nm/430 nm). D-E Heat-treated CsgA_High_ biofilms at different times and temperatures on 24-well plates. Images and absorbance spectra were taken and analyzed as mentioned before.

In order to test cell viability within biofilms after heat treatment, we performed a SYTO9/propidium iodide viability staining (**Figure 4A**). Before heat treatment 86.0 ± 12.1 % of the cells were alive, while viability after heat dropped to 11.9 ± 11.7 %. The heat treatment is sufficient to keep acidification to a minimum for a successful co-culture of ELMs and mammalian cells.

**Figure 4.**
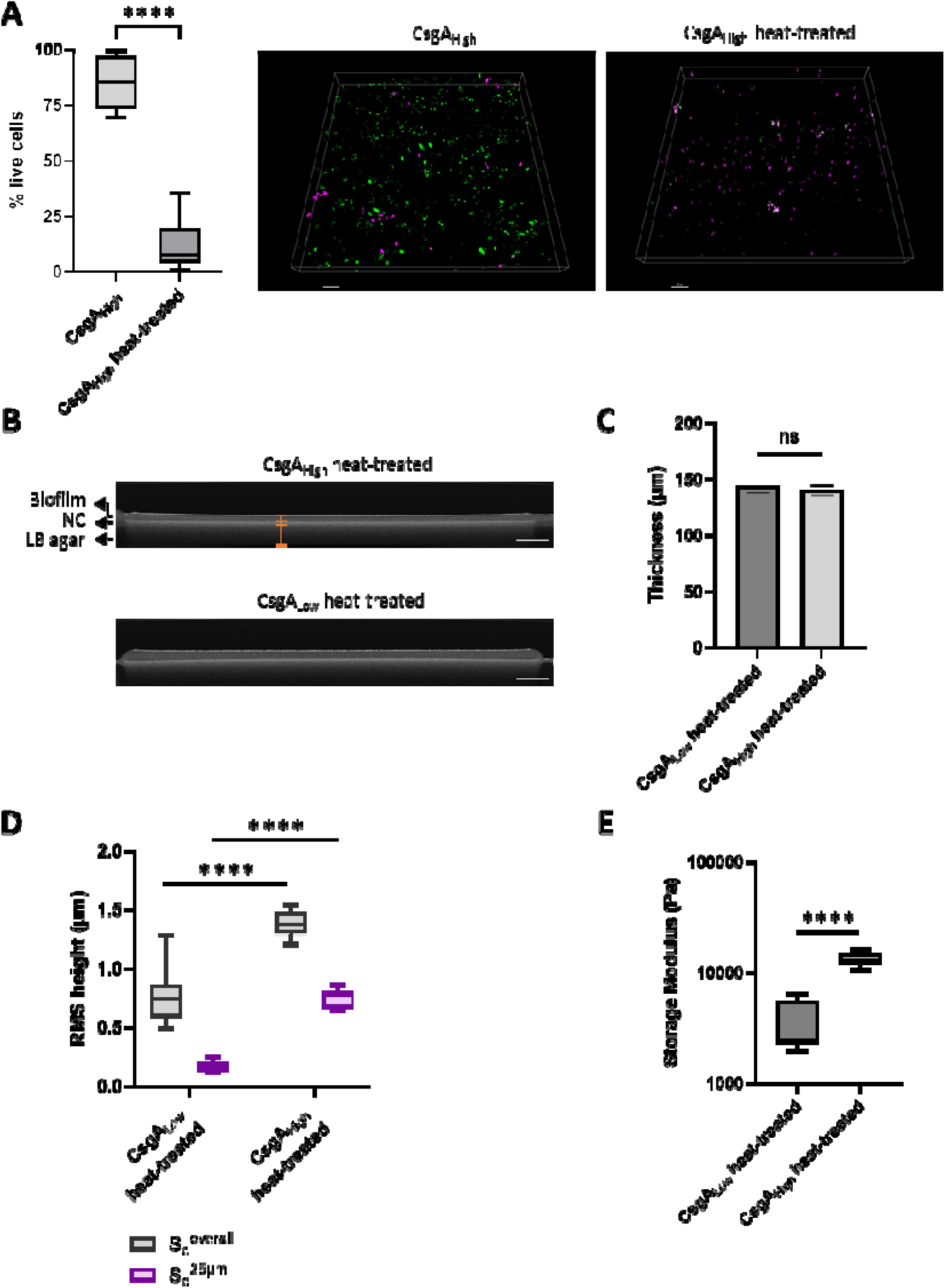
Analysis of heat-treated biofilms. A SYTO9/Propidium iodide viability staining of CsgA_High_ biofilms after heat treatment (65 °C for 1 h). The percentage of alive cells is displayed for non-treated and heat-treated biofilms (alive cells, green; dead cells, magenta). Scale bar: 10 µm. B Representative OCT cross-sectional image of biofilms heat-treated for 1 h at 65 °C. scale bar: 1000 µm. C Thickness of biofilms determined from the OCT cross-sectional images. A refractive index of 1.48 was used in the thickness calculations for both heat-treated biofilms (**Figure S2**). D Surface roughness of biofilms. S_q_ ^overall^ and S ^25^ ^µm^ were determined. E Rheological analysis of CsgA_Low_ and CsgA_High_ after heat treatment. Data in A and C-E are representative data from n = 3 replicates. Boxes represent median and upper and lower quartiles, whiskers represent min and max values. Mann-Whitney U test, **** *P* ≤ 0.0001, ns *P* > 0.05

We characterized the biofilms that were subjected to heat-treatment to evaluate if structural changes are introduced by this procedure. We measured the thickness, roughness and stiffness of biofilms heat-treated at 65 °C for 60 min. The thicknesses of CsgA_High_ heat-treated and CsgA_Low_ heat-treated biofilms were 140.3 ± 4.9 and 143.4 ± 5.9 µm, respectively (**Figure 4B** and **Figure 4C**). Further, thickness did not significantly change after heat-treatment compared to the untreated samples (*P* > 0.05, **Figure S7A**). Regarding roughness, the CsgA_High_ heat-treated biofilms exhibited higher S_q_ ^overall^ and S_q_ ^25µm^ values compared to the CsgA heat-treated biofilms, with S_q_ ^overall^ values of 1.39 ± 0.10 and 0.77 ± 0.21 µm and S_q_ ^25µm^ values of 0.75 ± 0.08 and 0.17 ± 0.04 µm, respectively (**Figure 4D**). This indicates that CsgA_High_ heat-treated has a rougher surface than CsgA_Low_ heat-treated biofilm. Interestingly, the roughness parameters for the biofilms decreased significantly following heat-treatment (*P* ≤ 0.01, **Figure S7B and S7C**), and this heating-induced decrease in roughness was stronger for the CsgA_Low_ biofilms.

Finally, regarding the stiffness of heat-treated biofilms, the difference between CsgA_High_ heat-treated and CsgA_Low_ heat-treated remains statistically highly significant (*P* ≤ 0.0001), with the CsgA_High_ exhibiting a storage modulus of 13.6 ± 1.7 kPa and CsgA_Low_ of 3.5 ± 1.7 kPa, maintaining a 4-fold difference (**Figure 4E**). The stiffness slightly decreases after heat treatment by 11.5 % and 13.1% for the CsgA_Low_ and CsgA_High_, respectively (*P* ≤ 0.05, **Figure S7D**). In addition to cell death, heat-treatment may cause protein denaturation, which together may alter the extracellular properties of biofilms.

Prior to examining the mechanosignaling response of HeLa cells to biofilms, we first calibrated the behavior of hYAP1 in HeLa cells seeded on well characterized PEG hydrogels with different stiffnesses. The soft hydrogels had a storage modulus of 0.09 ± 0.02 kPa and the stiff hydrogels of 9.7 ± 0.9 kPa (**Figure S8**). After cultivating cells on top of the hydrogels for 24 h, the cells were immunostained against hYAP1 and analyzed using a microscope (**Figure S9**). It was seen that hYAP1 localized to the nucleus on the stiff gel while localizing more to the cytosol on the soft gel. This is in line with findings of different studies describing the localization of YAP depending on substrate stiffness.[41–43]

As a next step, we performed immunostaining of HeLa cells seeded on the biofilms for 2 h or 6 h (**Figure 5A**). We observed that the cytoplasmic localization of hYAP1 is higher on CsgA_Low_ biofilms compared to cultivation on the CsgA_High_ ones. To quantitively analyze the difference between cytosolic and nuclear localization, we measured the fluorescence intensities of each and divided cytosolic by nuclear intensity (cyt/nuc) (**Figure 5B**). Analysis shows a significantly (*P* ≤ 0.0001) lower cyt/nuc ratio for CsgA_High_ than for CsgA_Low_ after 2 h and 6 h of cultivation. For further analysis, the areas of the whole cells were compared, since studies have shown that cells tend to spread more on stiffer surfaces (**Figure 5C**).[44] The cells seeded on CsgA_High_ biofilms, had significantly (*P* ≤ 0.0001) larger areas compared to the cells seeded on CsgA_Low_. This further underlines the hypothesis that cells sense the mechanical properties of biofilms.

**Figure 5.**
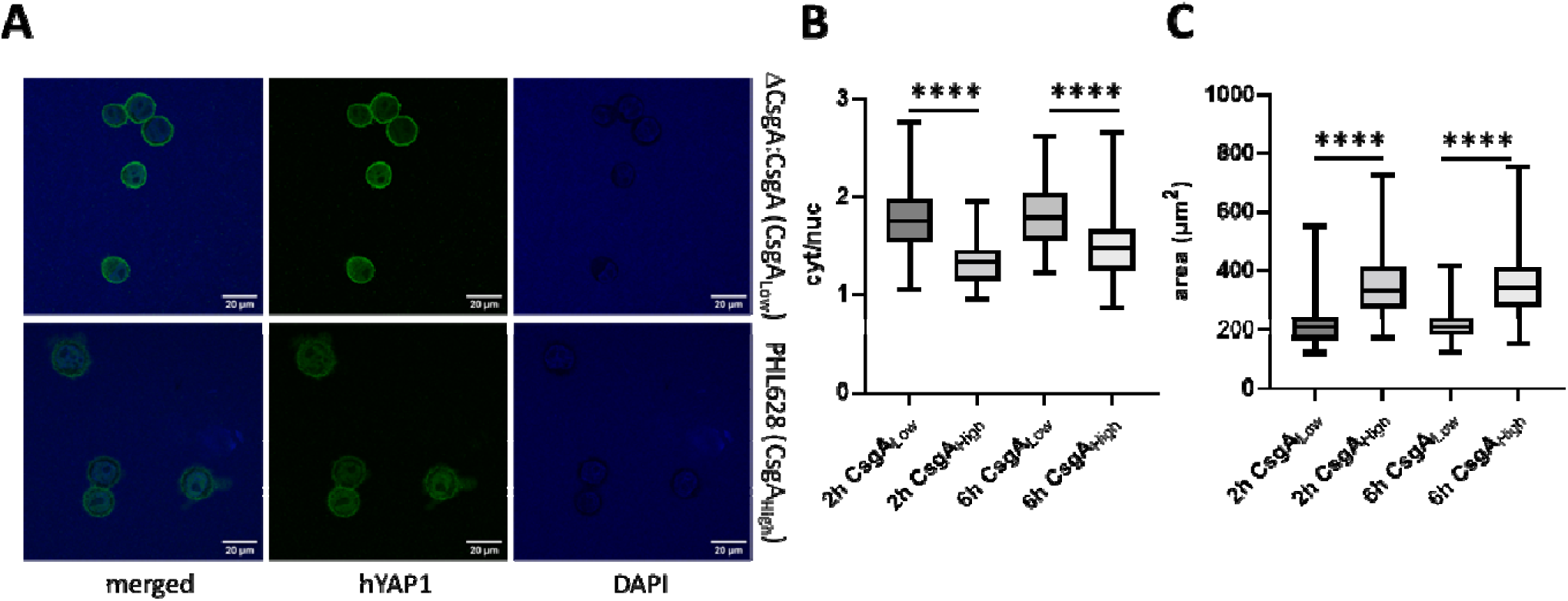
Immunostaining against hYAP1. A Representative microscopy images of hYAP1 stained HeLa cells cultivated for 6 h on two different biofilms CsgA_High_ and CsgA_Low_. The biofilms were grown for four days on nitrocellulose, heat treated for 1 h at 65 °C and then HeLa cells were incubated on top of the biofilms for 2 h or 6 h. B Quantitative analysis of cytosolic/nuclear (cyt/nuc) localization of hYAP1. The measured fluorescence intensity of cytosol and nucleus were normalized to the area and subsequently divided and plotted as box plots. C Areas of the cells plated on the different biofilms. Data in B-C are representative data from n = 3 replicates. Experiments were performed in biological duplicates and at least four randomly distributed images were taken from each sample. Boxes represent median and upper and lower quartiles, whiskers represent min and max values. Mann-Whitney U test, \*\*\*\**P* ≤ 0.0001

In order to expedite the assessment of mechanosignaling and avoid potential immunostaining artefacts, we developed a stable cell line that eliminates the need of immunostaining and allows the direct assessment of hYAP1 localization via fluorescence microscopy. To this end, HeLa cells were transduced via lentiviral particles encoding hYAP1 and H2B fused to the fluorescent proteins eGFP and BFP, respectively. H2B is a histone component and enables the visual identification of the nucleus, thereby facilitating the quantification of hYAP1 localization (**Figure S10**). Cells stably carrying both expression constructs were sorted by FACS and used for the following experiments.

We seeded the reporter cell line onto the biofilms and subsequently assessed hYAP1 localization and cell area. As previously observed in the immunostaining results, we saw preferred localization of hYAP1 to the cytosol when seeded on the softer biofilms (**Figure 6A**). Quantitative analyses further confirmed a significantly lower cyt/nuc ratio of hYAP1 and a significantly higher cell area for CsgA_High_ than for CsgA_Low_. (**Figure 6B** and **Figure 6C**).

**Figure 6.**
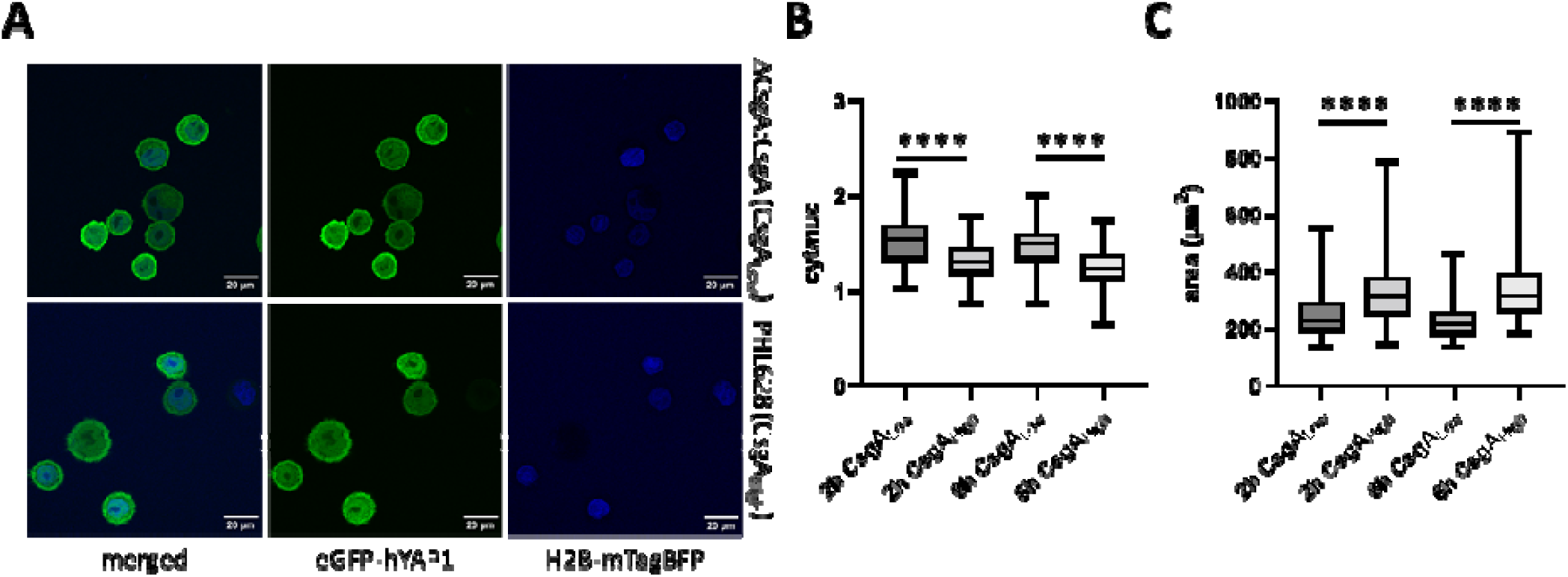
Analysis of hYAP1 localization using the reporter cell line on biofilms. A Representative microscopy images of stably transduced HeLa cells cultivated for 6 h on CsgA_High_ and CsgA_Low_ biofilms. The biofilms were grown for four days on nitrocellulose, heat treated for 1 h at 65 °C and then eGFP-hYAP1/H2B-mTagBFP transduced HeLa cells were incubated on top of the biofilms for 2 h or 6 h. B-C Analysis of cytosolic to nuclear localization of hYAP1 (cyt/nuc) and overall area of the cells. Data in B-C are representative data from n = 3 replicates. Experiments were performed in biological duplicates and at least four randomly distributed images were taken from each sample. Boxes represent median and upper and lower quartiles, whiskers represent min and max values. Mann-Whitney U test, \*\*\*\**P* ≤ 0.0001

With these experiments, we demonstrate that mammalian cells exhibit varying hYAP1 localization when cultivated on biofilms with different mechanical properties. On the softer engineered biofilms CsgA_Low_ cells show rather cytosolic localization of YAP, while on the stiffer CsgA_High_ biofilms cells display an enhanced nuclear localization. Previous studies have shown that, in addition to stiffness, surface roughness is another parameter that may influence mechanoresponse of mammalian cells.[45,46] Recent findings indicate that an increase in surface roughness from S_q_^overall^ = 0.054 µm to S_q_^overall^ = 0.880 µm, comparable to the surface roughness observed in our biofilms, has a greater impact on YAP nuclear localization in softer hydrogels (G’ = 3.8 kPa) than in stiffer hydrogels (G’ = 31.3 kPa); in the latter case, higher roughness may even reduce YAP nuclear localization.[46] Given these findings, we propose that the observed differences in YAP localization between the softer, smoother biofilm (CsgA_Low_, G’ = 3.5 kPa, S_q_^overall^ = 0.768 µm) and the stiffer, rougher biofilm (CsgA_High_, G’ = 13.6 kPa, S_q_^overall^= 1.389 µm) are primarily driven by variations in substrate stiffness. Progress in the control of biofilm mechanics may allow us to disentangle the effects of stiffness and roughness in future studies. Acting as a transcriptional activator of a large variety of genes, a dysregulation of YAP signaling is associated with a change in cell fate and function and has been correlated to different diseases.[47–49] For this reason, the mechanical properties of ELMs should be carefully tailored to match the mechanical properties of the cellular environment to avoid deregulating undesirable pathways. For example a mesenchymal stem cell sensing different mechanical input and an activation of YAP signaling might differentiate into an osteocyte, while in a cardiac progenitor cell it might lead to proliferation and differentiation into a cardiomyocyte.[50,51] In the skin alone, YAP localization differs between the cell types. A study showed that in the basal layer, YAP was found to be localized more in the nucleus, playing a role in skin homeostasis by promoting cell proliferation of stem and progenitor cells.[52] Inversely, in apical epithelial cells YAP is localized in the cytoplasm.

Further, YAP has been shown to be involved in fibrotic tissue formation in the lung, liver, heart and kidney. During fibrosis, fibroblasts differentiate, stimulated by YAP, into myofibroblasts, which produce various components of the extracellular matrix like collagens.[53] This leads to pathological hardening of the organ tissue which will subsequently lead to reduced organ function.[54] Especially, when conceiving ELMs for implantation, the mechanical properties should be well adjusted to avoid fibrosis formation and the subsequent encapsulation of the ELM which might limit its further (therapeutic) function.

## Conclusion

Engineered living materials (ELMs) are a new class of smart biomaterials, that found applications in a broad spectrum of disciplines. Increasingly interesting for biomedical applications, more and more ELMs are being developed focusing on their use as drug depots, wound dressings or biosensors. In this work, we demonstrate that HeLa cells respond to the presence of mechanically distinct biofilms via mechanosignaling. We believe that this effect is of high importance in therapeutic applications, since the mechanical properties of the therapeutic material could potentially affect surrounding cells in unforeseen ways. On the other hand, deliberately utilizing ELMs with different mechanical properties might facilitate the use of such materials as extracellular matrices in order to control cellular functions and developments. By applying heat treatments to limit bacterial growth, we could successfully show that the co-culture between biofilms and mammalian cells is feasible, further underlining the possibility of a broader use in cell culture applications. Overall, based on the observations from this study, the mere physical influence of a foreign material merits further investigation to better understand mechanotransductional influences of ELMs, while also considering future applications based on this effect.

## Experimental Methods

### Cloning

Plasmid pOK018 was assembled by Gibson cloning. The gene encoding CsgA (kindly provided by Dr. Neel S. Joshi, Northeastern University, Boston, MA) and mCherry were cloned into the two MCSs of pCDFDuet^TM^-1 (Novagen (EMD Millipore)) and the promotors were changed to the constitutive trc promotor.[24] For lentiviral plasmids pOK046 and pOK050, the backbone (pCDH_EF1a (Addgene plasmid # 72266) or pLVX-Puro (Clontech (TaKaRa))) was enzymatically digested. The inserts were assembled using fusion PCR, digested and cloned into the lentiviral backbone. All sequences of the open reading frames were verified.

### Biofilm Production

Biofilm producing *E. coli* cells (PHL628 or PHL628 ΔCsgA:pOK018) were grown from a glycerol stock o/n in LB Medium (10 g L^-1^ tryptone, 5 g L^-1^ yeast extract, 10 g L^-1^ NaCl, Roth) containing the respective antibiotics (PHL628: 40 µg mL^-1^ kanamycin, PHL628 ΔCsgA:pOK018: 40 µg mL^-1^ kanamycin, 36 µg mL^-1^ chloramphenicol, 50 µg mL^-1^ streptomycin) at 37 °C and 150 rpm.[35] The OD_600_ was measured and adjusted to the same value for all samples. Cells were subsequently plated on a nitrocellulose membrane (Roth) placed on top of an LB Agar plate without salt supplemented with kanamycin (10 g L^-1^ tryptone, 5 g L^-1^ yeast extract, 15 g L^-1^ agar, 40 µg mL^-1^ kanamycin) and grown for four days at room temperature.

### Mammalian Cell Culture and Development of Reporter Cell

For the production of lentiviral particles for transduction, HEK-293T cells were cultured in cell culture medium (DMEM supplemented with 10 % FCS, 100 U mL^−1^ penicillin and 100 µg mL^−1^ streptomycin) at 37 °C and 5 % CO_2_. After one day, cells were transfected using PEI with the packaging plasmids pLTR-G, pCD/NL-BH* ΔΔΔ and a plasmid coding for eGFP-hYAP1 (pOK046; eGFP-hYAP1 was kindly provided by Prof. Dr. Gerd Walz) or H2B-mTagBFP2 (pOK050) in a mass ratio of 1:1:2.[55,56] After incubation for 6 h, medium was removed and changed advanced DMEM, supplemented with 2 % FCS, 2 µL mL^-1^ cholesterol (5 mM in Phosphate Buffered Saline (PBS)), 2 µL mL^-1^ egg lecithin (5 mM in EtOH) and 1x chemical defined lipid concentrate (Gibco). After incubation for 2 days, the supernatant containing the lentiviral particles was removed, filtered using a 0.45 µm CA-S filter and stored at −80 °C until further use. For transduction, HeLa cells were cultured in cell culture medium at 37 °C and 5 % CO_2_. Supernatant containing the lentiviral particles (pOK046) was added and cells were further cultivated for 3 days. Afterwards, cells were detached using trypsin, resuspended in FACS buffer (PBS supplemented with 2 % FCS) and analyzed for transduction efficiency using an Attune NxT Flow Cytometer with autosampler (Thermo Fisher Scientific). Cells were subsequently sorted for eGFP expression. The cells were further cultivated and subsequently transduced with lentiviral particles (pOK050) as before. The cells were then sorted for eGFP and mTagBFP2 double-positive populations using a FACS Aria Fusion.

### Hydrogel synthesis

Star-shaped (8-arm) 10 kDa MW PEG-Vinylsulfone (PEG-VS, SKU: A10033-1) was purchased from JenKem Technology USA. RGD motif containing peptide (Ac-GCGYGRGDSPG-NH2) and two crosslinking peptides (Ac-GCRDWALQRPYPNPPVPDRCG-NH2 and Ac-GCRDGAGKYVAPNPPVPDRCG-NH2) were custom synthesized by Caslo ApS (Technical University of Denmark). All reagents were dissolved in 0.3 M triethanolamine (TEA, pH 8, Carl Roth, Art. No. 6300.1) prior to use and stored at −80 °C. For hydrogel preparation, 25 % (w/v) PEG-VS was mixed with 0.3 M TEA and 10 mM RGD peptide in a VS:Peptide molar ratio of 32:1, vortexed briefly, and allowed to react via Michael-type addition for 5 min at room temperature. Afterwards, 100 mM crosslinking peptides (1:1 mixture of both peptides) were added in a 1.25 molar excess to total VS groups, the sample was vortexed briefly and 50 µl drops were transferred onto a hydrophobically coated (Sigmacote) glass slide. Final concentrations of PEG-VS, RGD and crosslinking peptides were adjusted to achieve different stiffness (soft: 400 µM RGD, 8 mM crosslinking peptides, 1.6 mM PEG-VS; stiff: 1750 µM RGD, 35 mM crosslinking peptides, 7 mM PEG-VS). Generation of disc-shaped gels was achieved by placing two spacers with 0.8 mm height adjacent to the gels and placing a 3-aminopropyltriethoxysilane (APTES)-treated coverslip on top. Gels were allowed to polymerize for 1 hour at RT before they were transferred into a 6-well plate with 3 ml PBS o/n at 4 °C. The next day, gels were washed again with 3 ml PBS and subsequently equilibrated with 3 ml FCS-free cell culture medium (DMEM supplemented with 100 U mL^−1^ penicillin and 100 µg mL^−1^ streptomycin) for 4 h at 37 °C.

### Cell culture and immunostaining on hydrogels

HeLa cells were cultivated in cell culture medium prior to use and removed from the plate by trypsin treatment. Cells were seeded on top of the hydrogel placed in a 6-well plate at a density of 20,000 cells cm^-2^ and incubated for 24 h at 37 °C, 5 % CO_2_. Subsequently, immunostaining against YAP was performed as previously described.[57] In brief, cells on the hydrogels were fixed in 4 % paraformaldehyde (PFA) for 1 hour at RT. After washing with PBS, the samples were incubated with a 50 mM NH**_4_**Cl solution in PBS for 1 hour at RT to quench free aldehyde groups. Cells were subsequently permeabilized by 0.2 % Triton X-100 in PBS for 15 minutes at RT, followed by incubation in YAP blocking buffer (1 % (w/v) BSA, 5 % (v/v) fetal calf serum in PBS) for 1 hour. In between the permeabilization and blocking steps, samples were washed at least two times with PBS. The cells were then incubated with anti-YAP antibody (Santa Cruz Biotechnology, Dallas, TX, cat. no. sc-101199, dilution 1:50 in YAP blocking buffer) for 2 hours at RT. After 2 washing steps with PBS and another one with blocking buffer, the samples were incubated with an Alexa Fluor 488-conjugated donkey anti-mouse antibody (Thermo Fisher Scientific, cat. no. A-21202, dilution 1:250 in YAP blocking buffer) for 1 h. Subsequently the cells were washed 2 times in PBS and DAPI stained for 10 min at RT.

### Characterization of biofilms on Congo Red agar plates

Congo Red (CR) staining was performed as described by Zhou et al.[58] CR and Brilliant Blue were prepared at a stock concentration of 10 mg mL^-1^ in water, filter sterilized and stored at 4 °C. LB Agar plates without salt (10 g L^-1^ tryptone, 5 g L^-1^ yeast extract, 15 g L^-1^ agar) were supplemented with antibiotics as before, and with a final concentration of 50 µg mL^-1^ CR and 1 µg mL^-1^ Brilliant Blue. Biofilms were plated on top of nitrocellulose membranes as mentioned before and CR staining was observed after 4 days of culturing at RT.

### Spin-Down quantitative Congo Red binding assay

The protocol was adapted from previously published protocols.[59–61] The biofilms were resuspended in PBS and pelleted at 6,000 x g for 1 min. The pellet was resuspended in 0.075 mM Congo Red (CR, Sigma-Aldrich) in PBS and incubated at room temperature for 15 min. Subsequently, the mixture was centrifuge at 15,000 x g for 10 min and the absorbance of CR in the supernatant was measure at 490 nm in a Tecan Spark multimode microplate reader (Tecan Trading AG, Switzerland). Normalized curli fiber production was calculated by subtracting the absorbance obtained for the biofilms from that obtained for the 0.075 mM CR solution and normalized by the OD_600nm_ of the resuspended biofilm. Triplicates for each biofilm were measured.

### Determination of the thickness of biofilms

Biofilms grown on agar plates for 48 h at 25 °C were imaged using a Ganymede GAN312 OCT (Thorlabs Ganymede, Newton, NJ, USA) with a central wavelength beam of 880 nm and a Thorlabs LSM03 scanning objective. Images were acquired using the OCT software (ThorImage OCT 5.6) with the following parameters: an average of 8 for the A- and B- scans, an image frequency of 40 kHz for the A scan and the Hann Apodization window. The refractive index of biofilms was determined from OCT cross-sectional images employing a method equivalent to that proposed by Tearney et al. (1995).[62] In contrast to the authors, that divided the total thickness of a biofilm by its thickness above a planar reflector surface at a given region, we determined the refractive index by dividing the entire area of the biofilm by the area of the biofilm above the agar surface (Figure S2). The OCT images were analyzed using Fiji.[63] Triplicates were measured for both CsgA_High_ and CsgA_Low_ biofilms. The refractive index of non-treated and heat-treated biofilms was set to 1.37 and 1.48, respectively. These values correspond to the average refractive index of CsgA_High_ and CsgA_Low_ in each condition, non-treated and heat-treated.

### Surface roughness of biofilms

The roughness of biofilms was measured using a 3D confocal optical microscope (MarSurf CM explorer, Mahr, Göttingen, Germany). Each biofilm was measured at four different locations in 1.6 mm x 1.6 mm squares with a 10x/ NA = 0.3 air objective lens. Triplicates were measured for both CsgA_High_ and CsgA_Low_ biofilms. Data was processed with the MarSurf MfM Extended 9.2.9994 extended mountains software. First, the surface was leveled using the least square plane leveling method, then the removal of isolated outliers feature was applied, and finally the arithmetic mean height (Sa) and root mean square gradient (Sdq) were calculated.

### Live dead staining of biofilms

Heat-treated and non-heat-treated biofilms were placed on a µ-Dish 35 mm (Ibidi, Germany) and stained with a mixture of SYTO9 (Invitrogen by Thermo Fisher Scientific, USA) and PI (Invitrogen by Thermo Fisher Scientific, USA) at a final concentration of 5 µM and 30 µM in PBS, respectively, for 1 hour at RT. Heat treatment was carried out at 65 °C for 1 h. Samples were imaged using a Zeiss LSM 880 inverted confocal laser scanning microscope with Zen 2.3 SP1 software (Zeiss, Oberkochen, Germany). Images were acquired using a Zeiss Plan-Apochromat 63x/1.4 Oil DIC M27 objective with detection wavelengths 491–544 nm and 591–718 nm, and laser wavelengths of 488 and 543 nm respectively for live and dead bacterial populations. The Imaris Surface tool (Imaris v10.0, Bitplane, Zurich, Switzerland) was used to calculate the 3D volume of live and dead bacteria. Apart from cells in which only SYTO9 (live) or PI (dead) were detected, there were some in which both dyes were detected. These cells were classified as dead, since the presence of the PI dye in them indicated damage to the membrane. The percentage of live cells was calculated as the volume of live cells divided by the total volume of the live and dead cells.

### Rheological measurements

The rheological properties of hydrogels and biofilms were measured by small amplitude oscillatory measurements (SAOS) using a MCR302e rheometer (Anton Paar) with a parallel plate configuration (P-PTD200 and PP08/S, Anton Paar). Storage modulus G’ and loss modulus G’’ were recorded as dimensions for viscoelastic properties. Measurement parameters were individually optimized for disc shaped hydrogels and biofilms as described below.

Disc-shaped hydrogels were characterized at 37 °C with an initial normal force of 20 - 30 mN. To prevent dehydration, hydrogels were surrounded with cell culture medium.

Amplitude sweeps were performed with a constant frequency of 0.5 Hz in the range of 0.01 – 10 % deformation to determine the linear viscoelastic range (LVE). Frequency sweeps were performed in the range of 0.1 – 10 Hz with a constant deformation of 0.1 %. Subsequently, both moduli were recorded as time sweeps with 10 measuring points over 3 mins with a constant frequency of 0.5 Hz and deformation of 0.1 %. Biofilms were transferred onto the lower measurement plate on top of the nitrocellulose growth substrate and measured at room temperature with a constant normal force of 15 mN. A reduced steady force was required to maintain proper contact with the sample throughout the measurement without compromising structural integrity. Before measurement, 1 drop of PBS was added on top of the biofilm to prevent dehydration. Sweeps and measurements were performed with the parameters described above for the hydrogels, with the distinction that 1 Hz was used as constant frequency. For each hydrogel formulation and biofilm condition three replicate samples were recorded as time sweeps.

### Co-Culture of Mammalian Cells and Biofilms

For co-culturing, first, different antibiotics were tested. For this, biofilms were placed in 24-well plates and 500 µL antibiotic-free cell culture medium was added including different amounts of antibiotics (in µg mL^-1^, tetracycline: 10, 5, 2.5, 1.25, 0.6; erythromycin: 20, 10, 5, 2.5, 1.25; sulfamethoxazole: 10, 5, 2.5, 1.25, 0.6). Biofilms were incubated for 2 h at 37 °C and 5 % CO_2_ and the change of color of the medium was observed (from red to yellow). For this, photographic images were taken and 200 µL of the medium was further transferred to a flat bottom 96-well plate and the absorbance spectrum was measured using a SpectraMax iD5 (from 230-1000 nm in 10 nm steps). For analysis, peak absorbance at 430 nm (yellow) and 560 nm (red) was taken and compared between the different samples. To test heating for co-culturing, biofilms were placed (on agar plates) at RT, 50 °C, 60 °C and 65 °C for 30 and 60 minutes. Biofilms were incubated as before for 2 h, 4 h and 6 h and analysis was performed as before (images and spectra). For following experiments, biofilms were placed at 65 °C for 1 h for heat inactivation, unless otherwise stated. They were subsequently placed in 24-well plates for tissue culturing including the nitrocellulose membranes. HeLa cells (either CsgA_High_ or stable cell line) were detached using trypsin and 50,000 cells mL^-1^ were plated in antibiotic-free cell culture medium (DMEM, supplemented with 10 % FCS) on top of the biofilms at 37 C and 5 % CO_2_. After 2 to 6 h cells and biofilms were fixed using 4 % PFA for 1 h at RT. Samples with the stable cell line were subsequently washed with PBS and mounted on glass slides using Mowiol 4-88 (Roth). Samples with HeLa CsgA_High_ cells were further processed with immunostaining.

### Immunostaining

For immunostaining of HeLa cells seeded on biofilms, an adapted protocol was used. Differences include: no quenching of free aldehyde groups was performed, no washing steps between permeabilization and blocking was performed, no washing step with blocking buffer was included after staining with the first antibody in order to ensure stability of biofilms throughout the treatment.

### Microscopy and Analysis

Cells were imaged using a Zeiss LSM 880 inverted laser scanning confocal microscope equipped with Zeiss Plan-Apochromat 63x/1.4 Oil DIC M27 objective. For analysis, Fiji was used and images were processed manually.[63] First, nuclei (nuc) were defined as regions of interest (ROIs) in the DAPI/BFP channel and transferred onto the GFP channel. The corresponding cells (all) were defined as ROIs in the GFP channel and the GFP intensity (integrated density (IntDent)) was measured. For calculation of the cytosolic to nuclear (cyt/nuc) ratio of YAP localization, the following equations were used (**Equation 1, 2 and 3**).

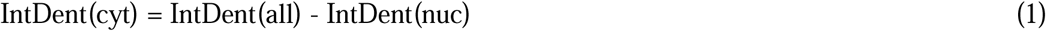

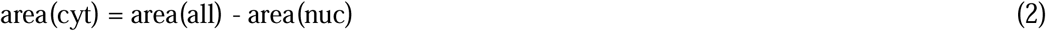

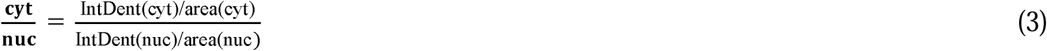

### Statistics and software

If not stated otherwise, the values indicated in the main text represent mean ± SD. For statistical analysis, single values were used and after normality tests, a Mann-Whitney U test was performed using GraphPad Prism version 9.2.0 for Windows, GraphPad Software, Boston, Massachusetts USA, www.graphpad.com. Figure 1 and ToC were created with BioRender.com.

## Supporting Information

Supporting Information is available from the Wiley Online Library or from the author.

## Acknowledgements

We thank the staff of the Life Imaging Center (LIC) in the Center for Biological Systems Analysis (ZBSA) of the University of Freiburg for help with microscopy resources, and the support in image recording and analysis. We acknowledge the technical assistance of the Signalling Factory Core Facility for help with flow cytometry and FACS sorting. We thank the Lighthouse Core Facility for their support with cell sorting (funded in part by the Medical Faculty, University of Freiburg (Project Numbers 2021/A2-Fol; 2021/B3-Fol) and the DFG (Project Number 450392965)). We thank Anthony Hay for providing the PHL628 and PHL628Δ*CsgA* bacterial strains. We acknowledge the Fluorescence Microscopy Facility at INM - Leibniz Institute for New Materials for support and advice with microscopy. This work was supported by the European Research Council (ERC) Project ID 101053857 – STEADY and the VolkswagenStiftung Experiment Program – 9A552.

Received: ((will be filled in by the editorial staff))

Revised: ((will be filled in by the editorial staff))

Published online: ((will be filled in by the editorial staff))

## Notes

### Competing Interest Statement

The authors have declared no competing interest.

