## Supplementary material for "Bacterial Engineered Living Materials modulate Mechanosignaling in Mammalian Cells": SupplemetaryInformation

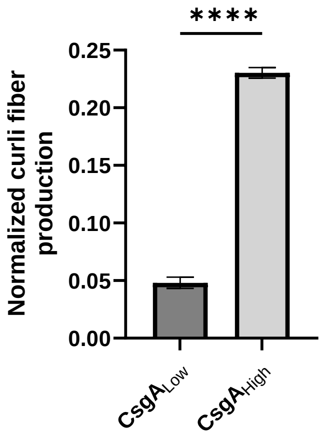


**Figure S1**. Quantification of curli fibers in biofilms grown from $CsgA_{High}$ and $CsgA_{Low}$ cells. Biofilms were grown on nitrocellulose membranes placed on LB agar plates for four days prior to performing a spin-down quantitative Congo Red (CR) binding assay. The normalized curli fiber production for each biofilm was obtained by normalizing the CR absorbance at 490 nm to the $OD_{600nm}$ of each biofilm. Data is representative from n = 3 replicates. Bars represent mean ± SD. Mann-Whitney U test, ****$P \leq 0.0001$

**A**

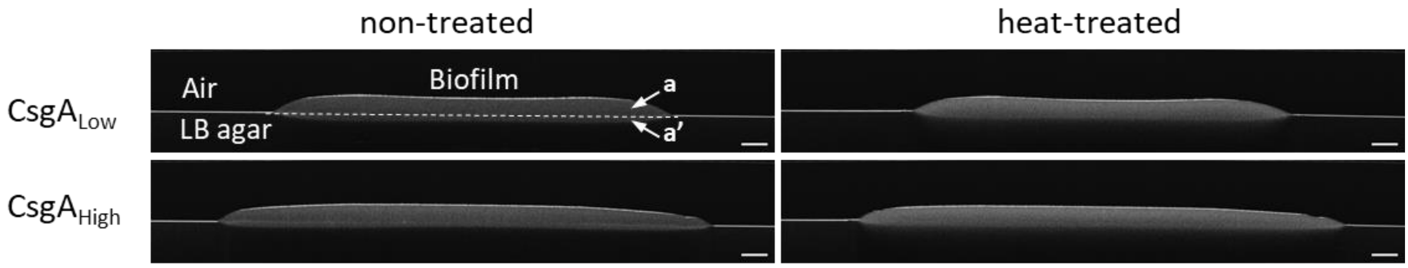


**B**

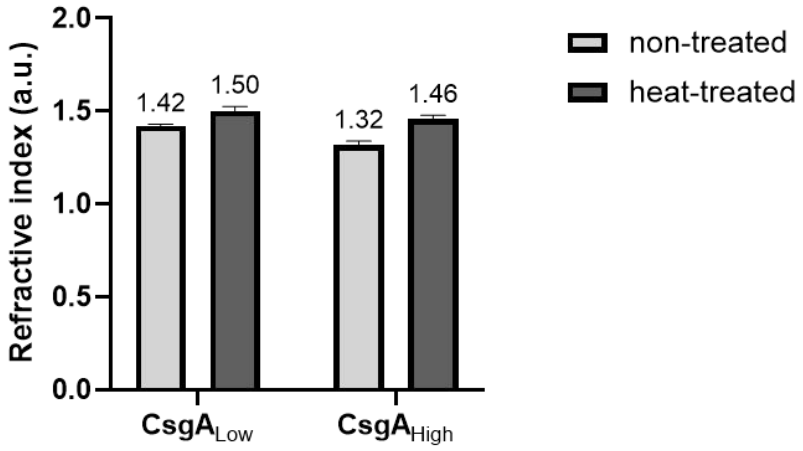


**Figure S2.** Determination of the refractive index of biofilms using optical coherence tomography (OCT) **A** Representative OCT images of biofilms grown on agar plates. To determine the refractive index of biofilms using OCT, biofilms were placed on a planar reflector surface. To this end, biofilms were grown on LB Agar plates for 4 days at room temperature. a: biofilm area above the LB agar surface, a': biofilm area under the LB agar surface. Scale bar: 500 µm **B** Refractive index determination of biofilms. The refractive index of biofilms was determined from the OCT images represented in part A by dividing the entire area of the biofilm (a + a ') by the area of the biofilm above the agar surface (a). The OCT images were analyzed using Fiji. Triplicates were measured for each condition. Bars represent mean ± SD.

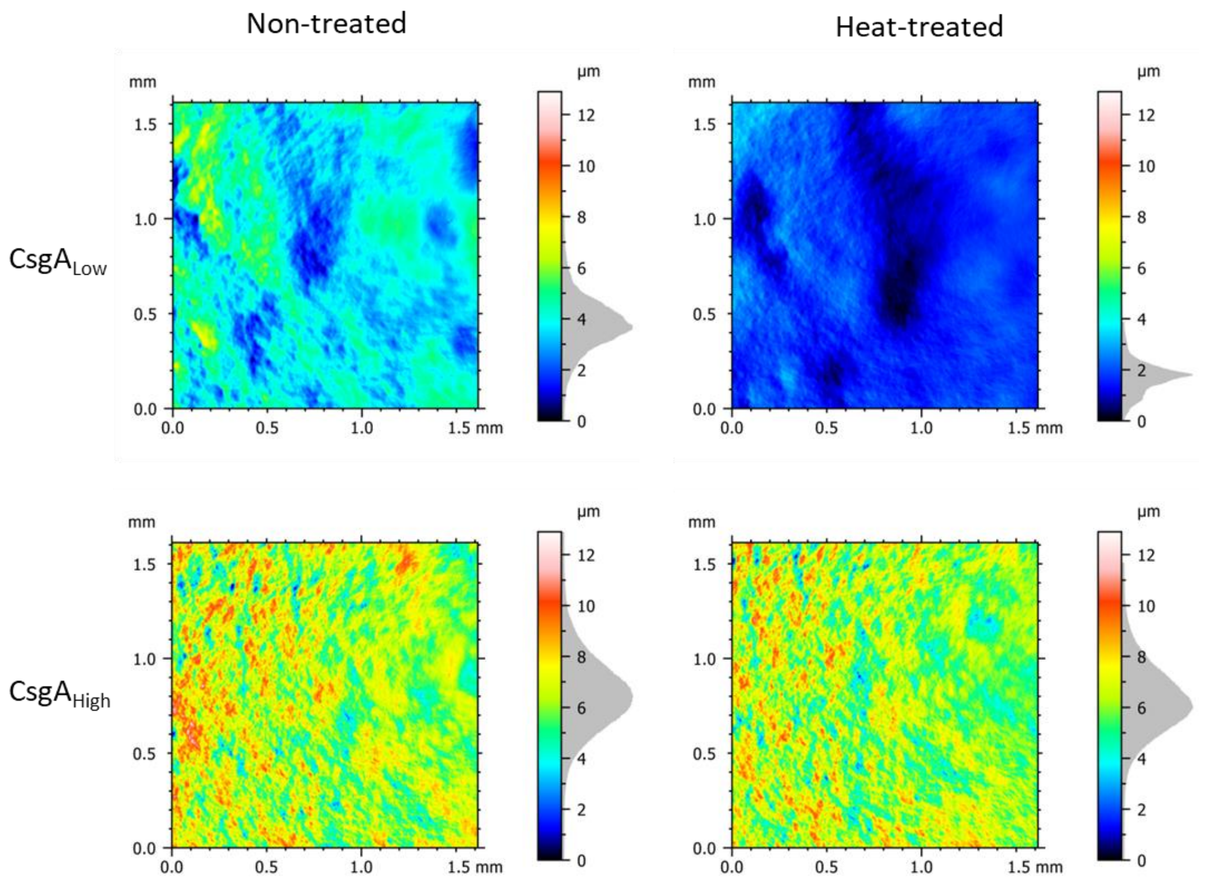


**Figure S3.** Surface analysis of biofilms. CsgA$_{Low}$ and CsgA$_{High}$ biofilms were grown for four days on nitrocellulose membranes placed on LB agar plates and heat treated at 65 °C for 60 min. Then the biofilms were analyzed by 3D confocal microscopy and the surface height was graphically represented. Exemplary images of the surface topography are shown.

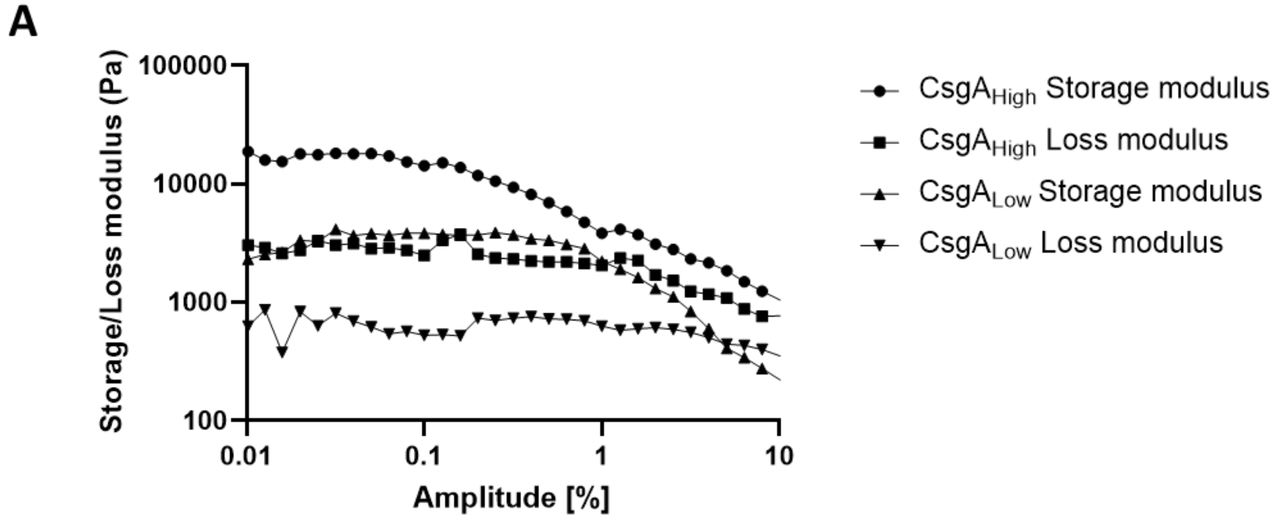


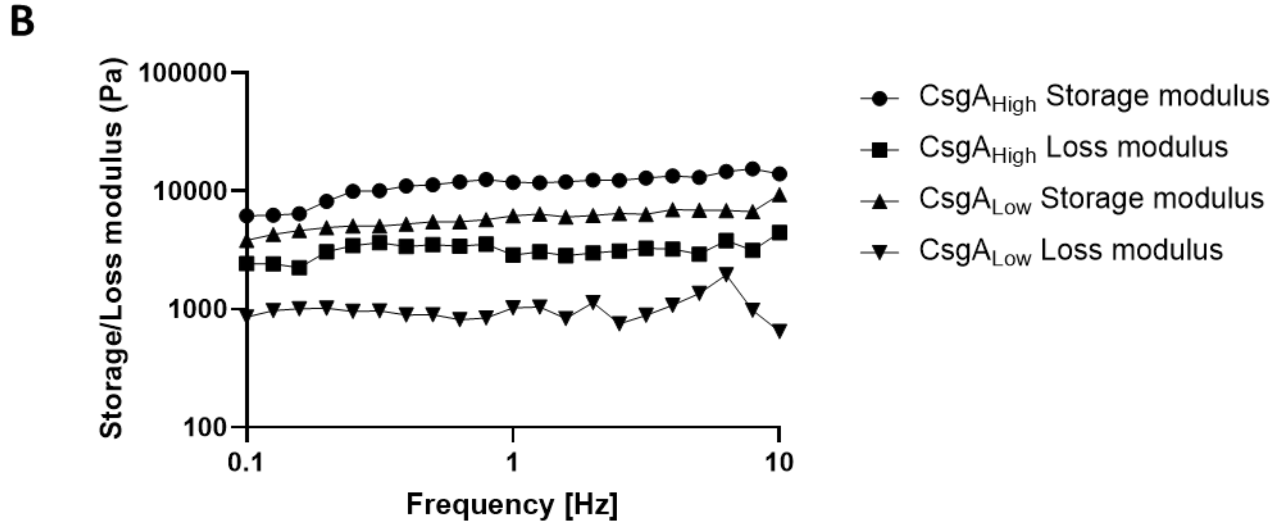


**Figure S4.** Rheological analysis of biofilms. $CsgA_{High}$ and $CsgA_{Low}$ biofilms were grown for four days and analyzed using a rheometer. **A** Amplitude sweep analysis. **B** frequency sweep analysis.

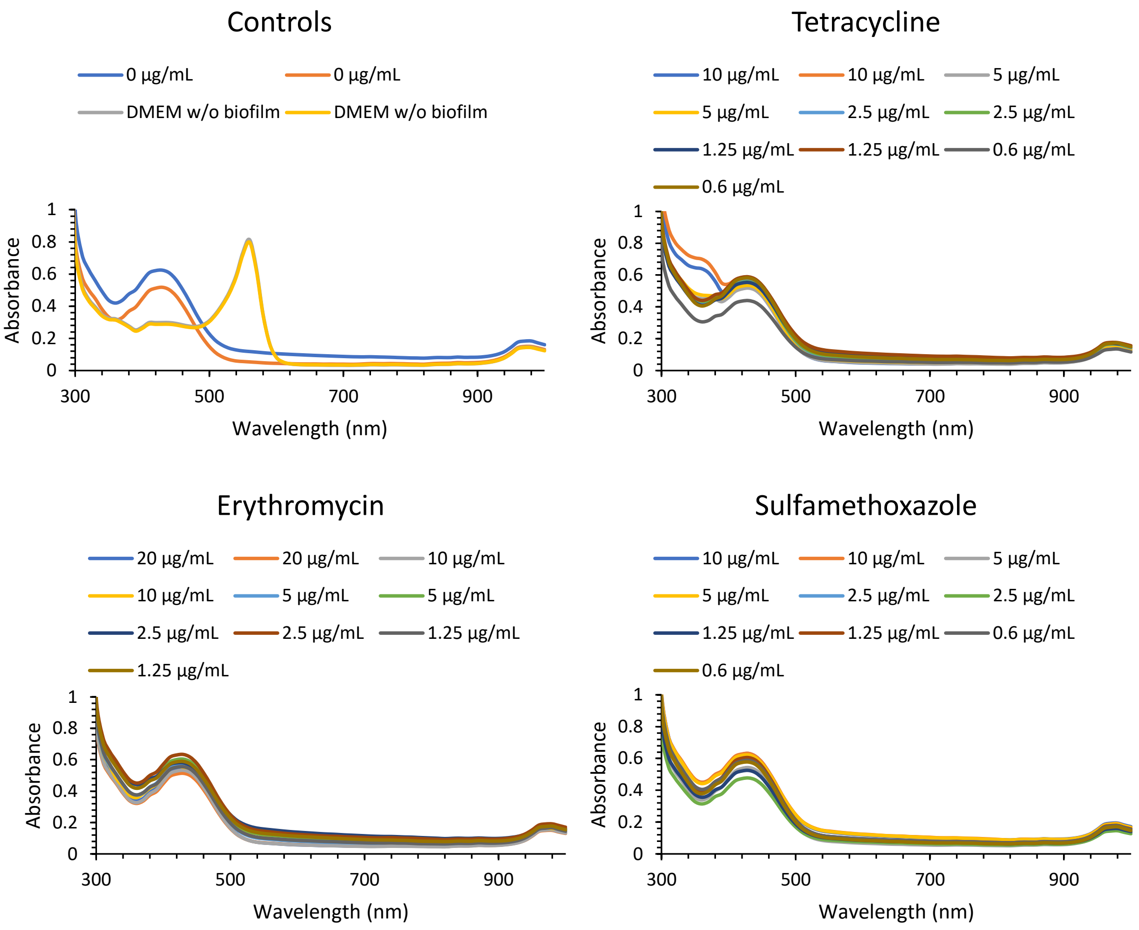


**Figure S5.** Absorption spectra measured after treatment with antibiotics. $CsgA_{High}$biofilms were grown for four days on nitrocellulose membranes placed on LB agar plates, subsequently placed in 24-well plates and incubated for 2 h in cell culture medium containing the indicated concentrations of antibiotics. The medium was taken and the absorbance spectrum was measured for each sample. Each experimental setup was performed in duplicates.

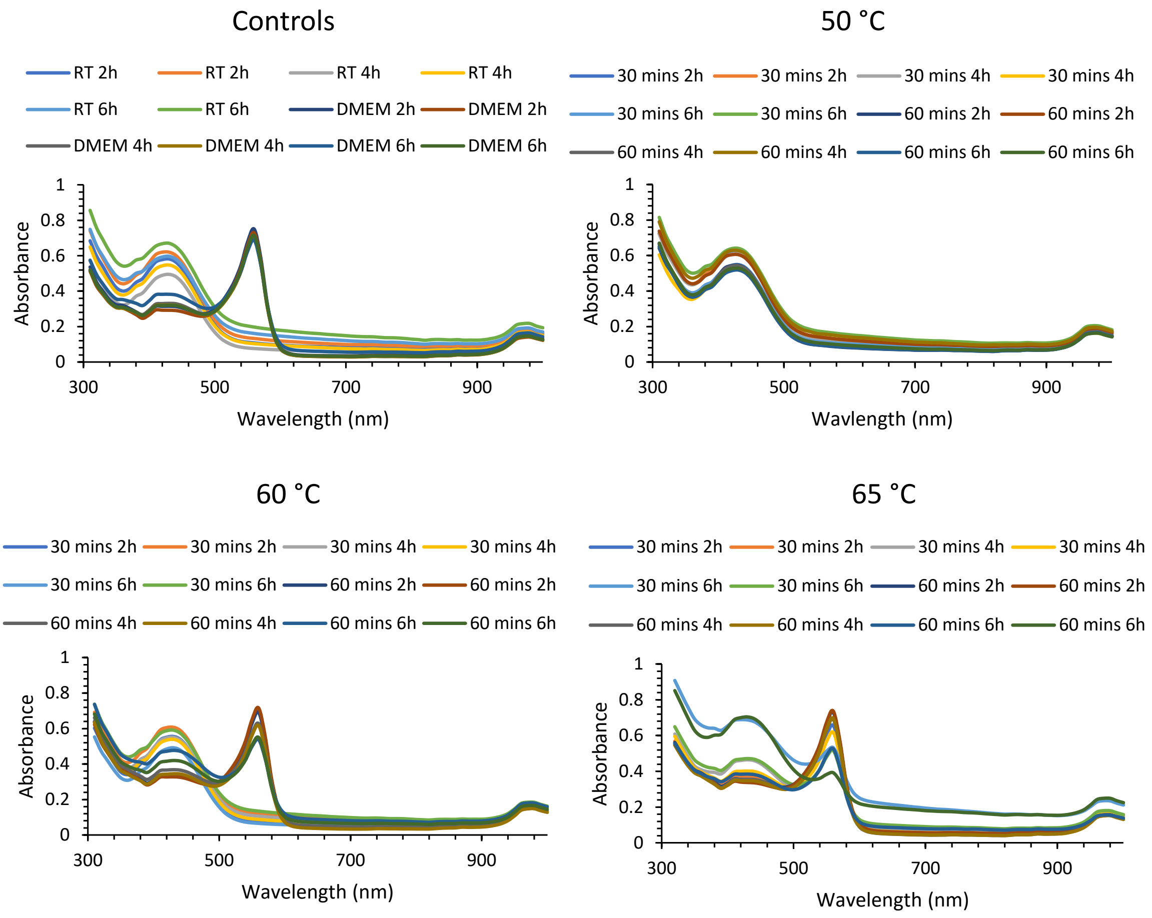


**Figure S6.** Absorption spectra measured after heat treatments. $CsgA_{High}$ biofilms were grown for four days on nitrocellulose membranes placed on LB agar plates and subsequently heat treated at indicated temperatures and times (30 minutes or 60 minutes). Biofilms were placed in 24-well plates and incubated for 2 h, 4 h or 6 h in cell culture medium. The medium was taken and the absorbance spectrum was measured for each sample. Each experiment set up was performed in duplicates.

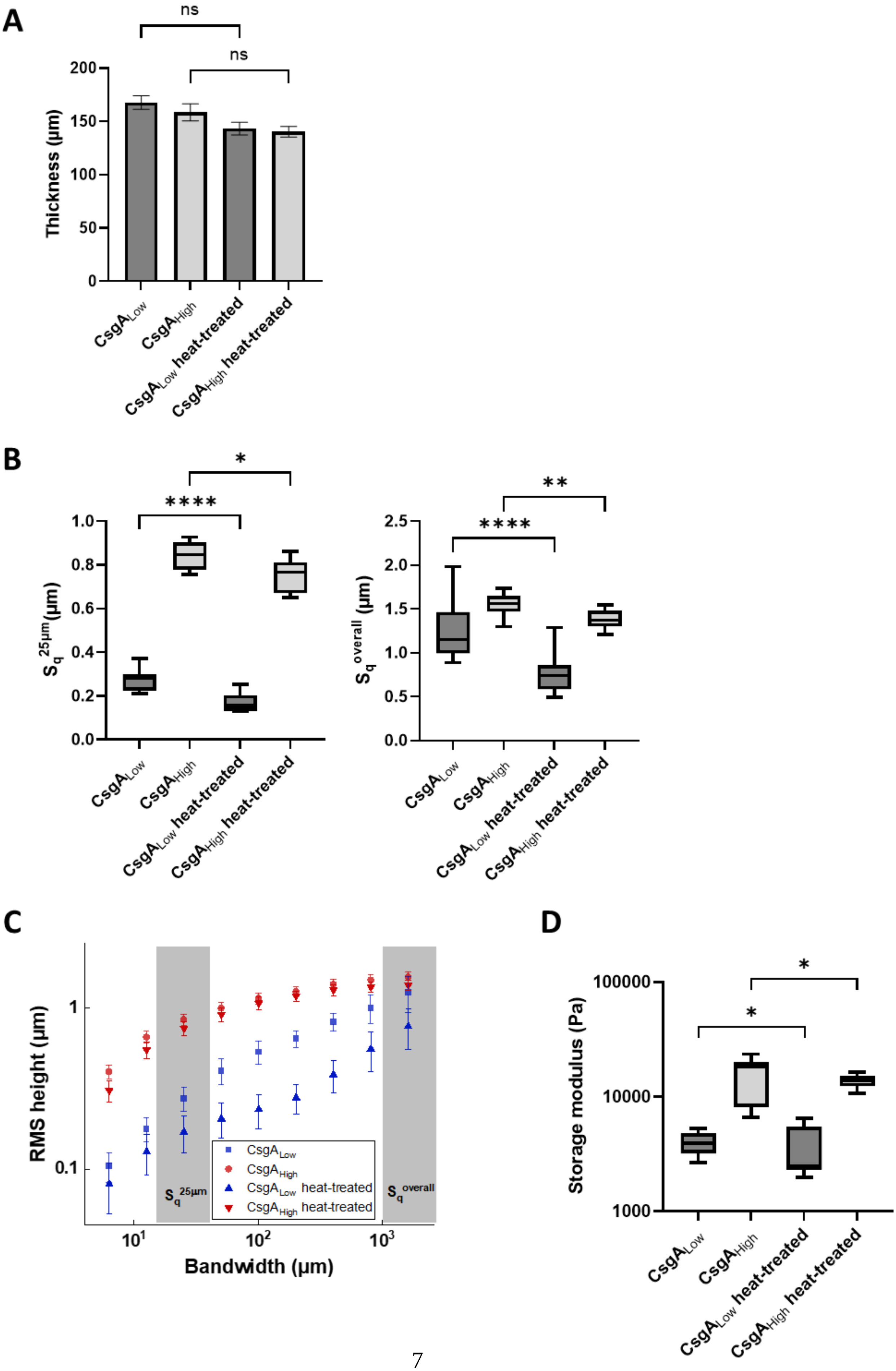
A
ns
ns
Thickness (µm)
200
150
100
50
0
CsgA_Low
CsgA_High
CsgA_Low heat-treated
CsgA_High heat-treated
B
*
****
1.0
0.8
0.6
0.4
0.2
0.0
S_q^25µm (µm)
**
****
2.5
2.0
1.5
1.0
0.5
0.0
S_q^overall (µm)
C
RMS height (µm)
1
0.1
S_q^25µm
S_q^overall
CsgA_Low
CsgA_High
CsgA_Low heat-treated
CsgA_High heat-treated
10^1
10^2
10^3
Bandwidth (µm)
D
*
*
Storage modulus (Pa)
100000
10000
1000

**Figure S7**. Comparison of biofilms before and after heat-treatment. **A** Thickness of biofilms before and after heat-treatment. Thickness of the biofilms was determined from optical coherence tomography (OCT) cross-sectional images. The refractive index for non-treated and heat-treated biofilms was set to 1.37 and 1.48, respectively (**Figure S2**). Heat-treatment was carried out at 65 °C for 60 min. **B** Surface roughness of biofilms before and after heat-treatment. Heat-treatment was performed at 65 °C for 1 h. The scale of overall roughness ($S_q^{overall}$) and the scale of a single mammalian cell (roughness $S_q^{25µm}$) are plotted. **C** The root-mean-square height deviation from the mean height (RMS height) is plotted as function of the spatial bandwidth, i.e. a characteristic length scale. The scale of overall roughness ($S_q^{overall}$) and the scale of a single mammalian cell (roughness $S_q^{25µm}$) are indicated. **D** Storage moduli of biofilms before and after heat-treatment. Heat-treatment was performed at 65 °C for 1h. The storage modulus significantly decreased after heat-treatment, for both $CsgA_{Low}$ and $CsgA_{High}$ biofilms. Data is representative from *n* = 3 replicates. Bars represent mean ± SD. Boxes represent median and upper and lower quartiles, whiskers represent min and max values. Mann-Whitney U test, **** $P \leq 0.0001$, ** $P \leq 0.001$, ** $P \leq 0.01$, * $P \leq 0.05$, ns $P > 0.05$

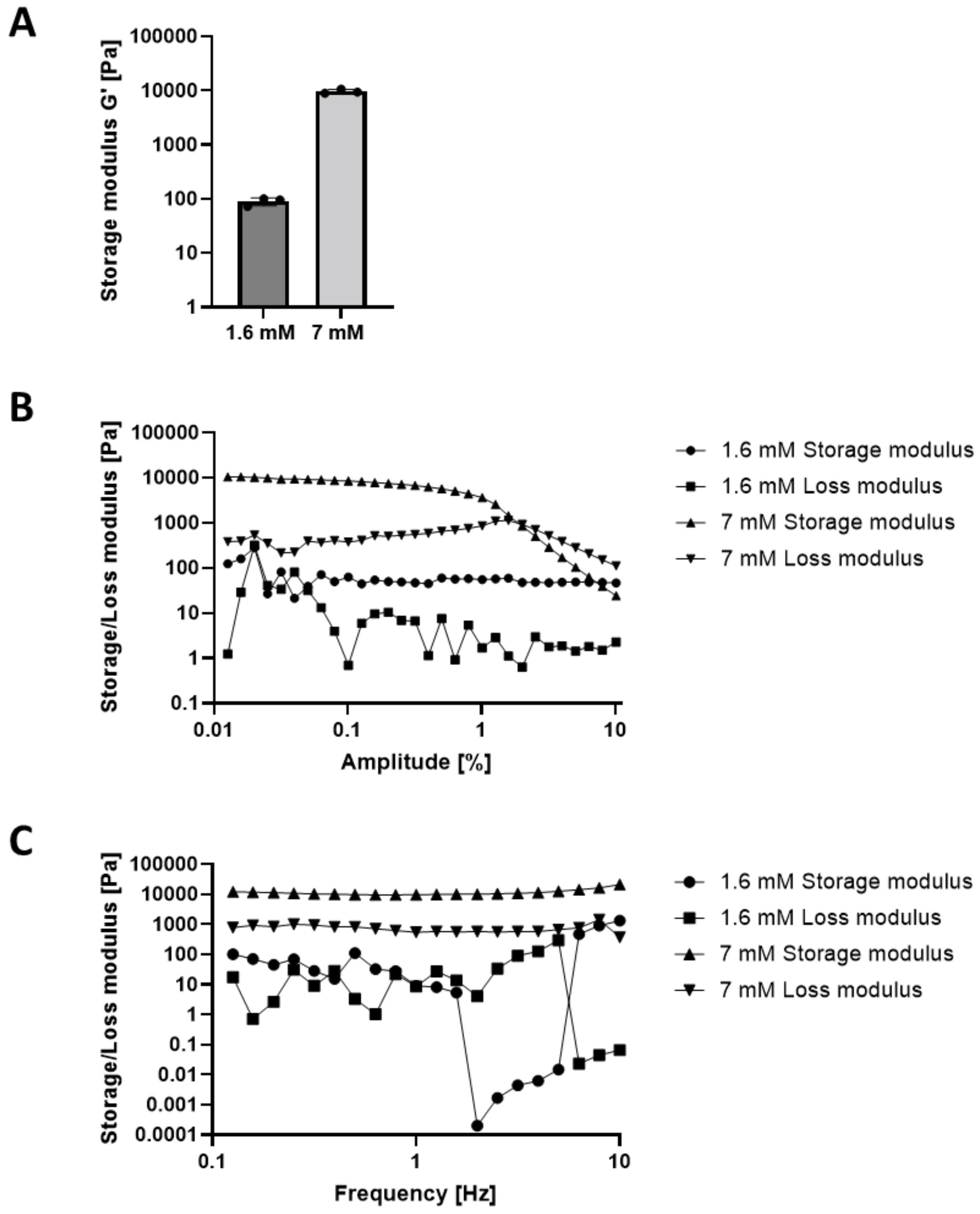


**Figure S8.** Characterization of hydrogel stiffness. **A** Hydrogels were prepared using different amounts of PEG-VS and the stiffness was measured using a rheometer. **B** Frequency sweeps of both hydrogels and **C** Amplitude sweeps of both hydrogels.

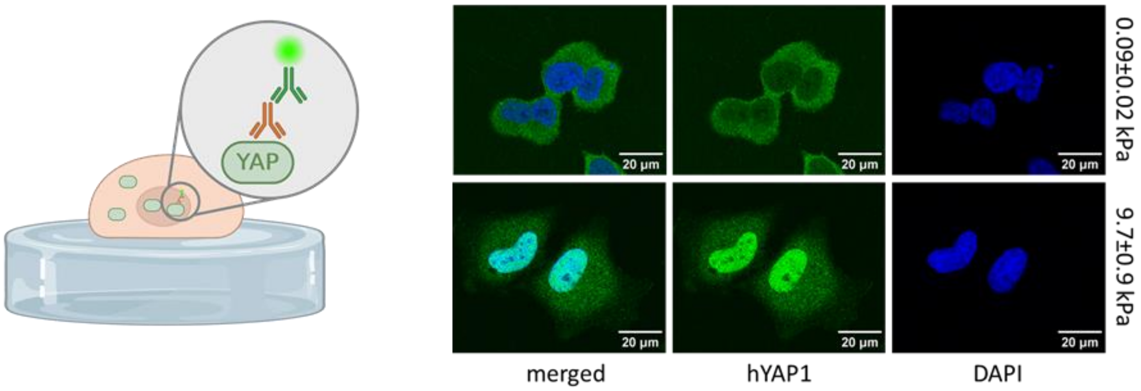


**Figure S9**. Immunostaining against hYAP1 on PEG gels. WT HeLa cells were seeded on PEG hydrogels with different storage moduli and subsequently immunostained against hYAP1.

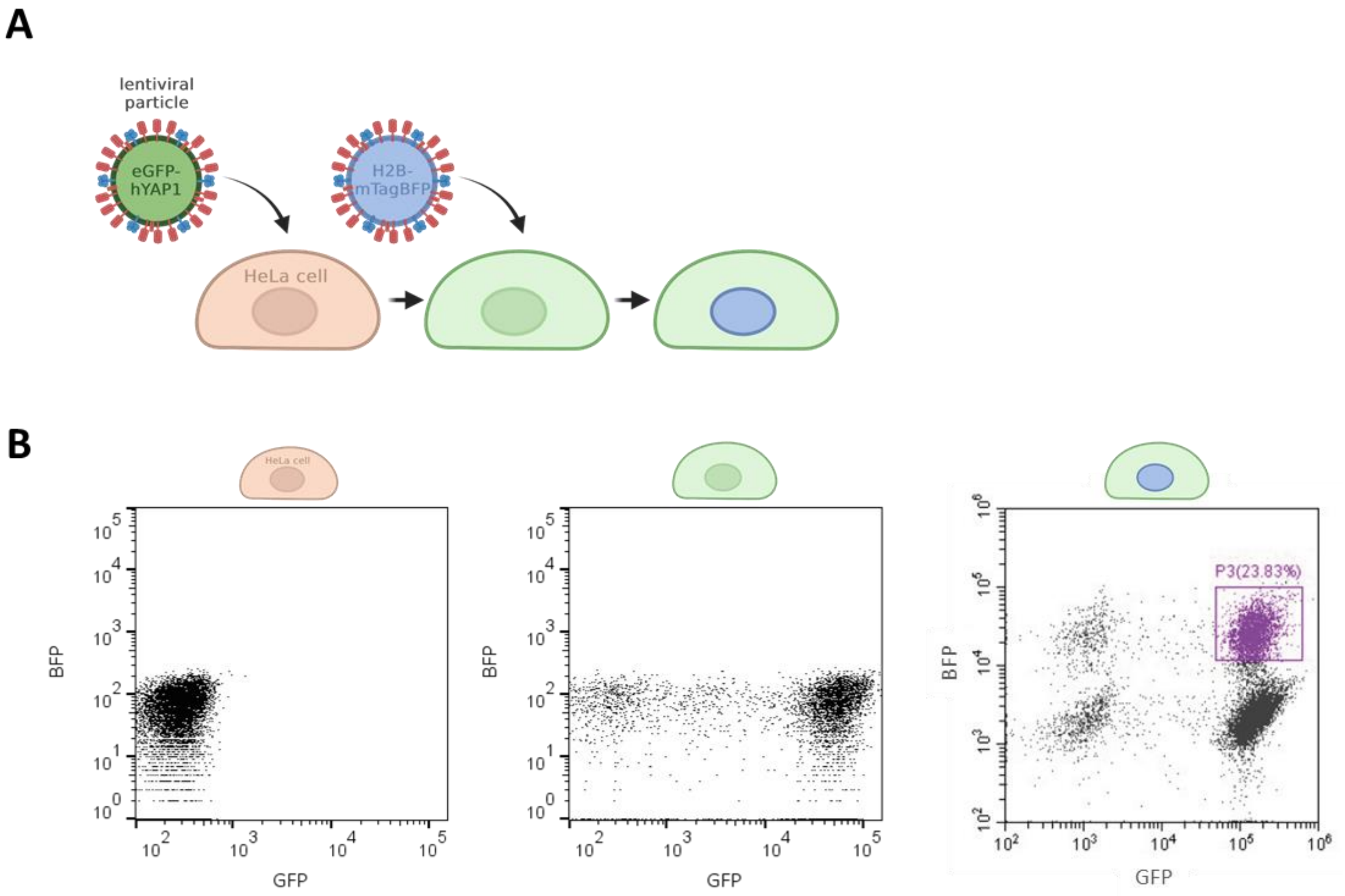


**Figure S10. A** Schematic overview of stable cell line generation by transduction using lentiviral particles. HeLa cells were first transduced with lentiviral vectors encoding eGFP-hYAP1 and subsequently with H2B-mTagBFP. **B** Flow cytometry data of non-transduced HeLa cells, transduced with eGFP-hYAP1 and with H2B-mTagBFP. Double positive cells marked with “P3”, were sorted and used as reporter cells further on.
